# Localization of PPM1H phosphatase tunes Parkinson’s disease-linked LRRK2 kinase-mediated Rab GTPase phosphorylation and ciliogenesis

**DOI:** 10.1101/2023.06.15.545139

**Authors:** Wondwossen M. Yeshaw, Ayan Adhikari, Claire Y. Chiang, Herschel S. Dhekne, Paulina S. Wawro, Suzanne R. Pfeffer

**Affiliations:** Department of Biochemistry, Stanford University School of Medicine, Stanford, CA 94305-5307 USA; Aligning Science Across Parkinson’s (ASAP) Collaborative Research Network, Chevy Chase, MD 20815

**Keywords:** LRRK2, Parkinson’s disease, Rab GTPase, protein phosphatase, amphipathic helix

## Abstract

PPM1H phosphatase reverses Parkinson’s disease-associated, LRRK2-mediated Rab GTPase phosphorylation. We show here that PPM1H relies on an N-terminal amphipathic helix for Golgi localization. The amphipathic helix enables PPM1H to bind to liposomes in vitro, and small, highly curved liposomes stimulate PPM1H activity. We artificially anchored PPM1H to the Golgi, mitochondria, or mother centriole. Our data show that regulation of Rab10 GTPase phosphorylation requires PPM1H access to Rab10 at or near the mother centriole. Moreover, poor co-localization of Rab12 explains in part why it is a poor substrate for PPM1H in cells but not in vitro. These data support a model in which localization drives PPM1H substrate selection and centriolar PPM1H is critical for regulation of Rab GTPase-regulated ciliogenesis. Moreover, Golgi localized PPM1H maintains active Rab GTPases on the Golgi to carry out their non-ciliogenesis-related functions in membrane trafficking.

**Significance Statement:** Pathogenic, hyperactive LRRK2 kinase is strongly linked to Parkinson’s disease and LRRK2 phosphorylates a subset of Rab GTPases that are master regulators of membrane trafficking. PPM1H phosphatase specifically dephosphorylates Rab8A and Rab10, the major LRRK2 substrates. Here we provide novel cell biological and biochemical insight related to the localization and activation of PPM1H phosphatase. Understanding how PPM1H modulates LRRK2 activity is of fundamental interest and also important, as activators of PPM1H may eventually benefit Parkinson’s disease patients.

## Introduction

Hyperactive, pathogenic mutations in Leucine Rich Repeat Kinase 2 (LRRK2) represent the most common cause of inherited Parkinson’s disease (1). Phosphoproteomics revealed that a subset of Rab GTPases is selected for LRRK2 action in cells (2,3), especially Rab10 and Rab8A. Rab phosphorylation occurs within the so-called Switch 2 region that is critical for Rab effector protein recognition, guanine nucleotide exchange factor activation, and GDI-mediated recycling (4). Thus, Rab phosphorylation blocks the abilities of certain Rab GTPases to be activated and to bind to many of their cognate effector proteins (2); it also entraps them on the compartment upon which they are phosphorylated because they cannot bind their recycling chaperone, GDI (2,5).

Despite loss of binding capacity to many effector proteins, phosphorylated Rab proteins gain phospho-specific binding capacity to a new set of proteins including RILPL1, RILPL2 and MyoVa and LRRK2 (3,6,7). In particular, binding of phosphoRab10 to RILPL1 is necessary and sufficient for LRRK2 blockade of primary cilia formation (8). LRRK2-generated phosphoRab10-RILPL1 and phosphoRab10-MyoVa complexes block primary cilia formation by a mechanism upstream of recruitment of TTBK2 kinase to the mother centriole (6, 9). Pathogenic LRRK2 also enhances cilia loss by a yet to be determined, Rab10 and RILPL1-independent pathway (9). Importantly, only a few percent of the total pool of Rab proteins is phosphorylated at steady state (2,10), yet this pool is sufficient to block cilia formation in a dominant manner (3,8).

Rab GTPase phosphorylation turns over extremely rapidly: treatment of cultured cells with LRRK2 inhibitors leads to complete dephosphorylation within just a few minutes (10). Elucidating how phosphatases control Rab phosphorylation is thus critical to our understanding the consequences of LRRK2-mediated Rab phosphorylation. Berndsen et al. (11) reported the discovery that PPM1H is a Rab-specific phosphatase that can reverse LRRK2-mediated Rab phosphorylation. Multiple lines of evidence confirm PPM1H’s role in phosphoRab biology. Loss of PPM1H phenocopies hyperactivation of LRRK2 in cell culture (11) and mouse brain (12). Moreover, after addition of LRRK2 inhibitors, cells lacking PPM1H recover Rab phosphorylation at about half the rate of control cells. Finally, unbiased mass spectrometry of proteins bound to a substrate trapping PPM1H mutant showed strong enrichment for Rab10, Rab8A and Rab35 (11).

We showed previously that exogenously expressed PPM1H is localized primarily to the Golgi complex, with additional localization to cytosolic and mitochondria-associated pools (11). Here we investigate the molecular basis for PPM1H Golgi localization and probe the importance of PPM1H localization on Rab10 and Rab12 phosphorylation and function.

## Results and Discussion

To identify the portion of PPM1H responsible for its Golgi localization, we aligned the PPM1H sequence with that of its nearest relative, PPM1J that is not Golgi localized (Fig. 1A,B,D). Amino acid insertions between PPM1H residues 115-133 and 205-218 distinguished the PPM1H sequence from that of PPM1J; similarly, PPM1J’s amino terminal region contains an insert between residues 13 and 48 in comparison with PPM1H (Fig. 1A). Analysis of mutant PPM1H proteins missing either or both of these loops revealed that both were dispensable for PPM1H co-localization with the Golgi marker, ACBD3 (Fig. 1C). In contrast, removal of the N-terminal 37 residues (Δ37) led to a cytosolic distribution of PPM1H protein (Fig. 1C, D). PPM1J was detected in small structures in the perinuclear region that lacked the cisternal appearance of Golgi-associated ACBD3 and may represent late endosomes (Fig. 1B). Co-staining of cells expressing PPM1H and PPM1J with the mitochondrial marker, mitofilin, confirmed that most of the proteins were not mitochondrially associated (Supplemental Figure 1A, B); nevertheless, a small proportion of PPM1H co-localized with mitofilin as we have reported previously (11). PPM1H has also been reported to be phosphorylated (13); non-phosporylatable S124A and S211A PPM1H proteins were not altered in their localizations (Supplemental Figure 2).

**Figure 1.**
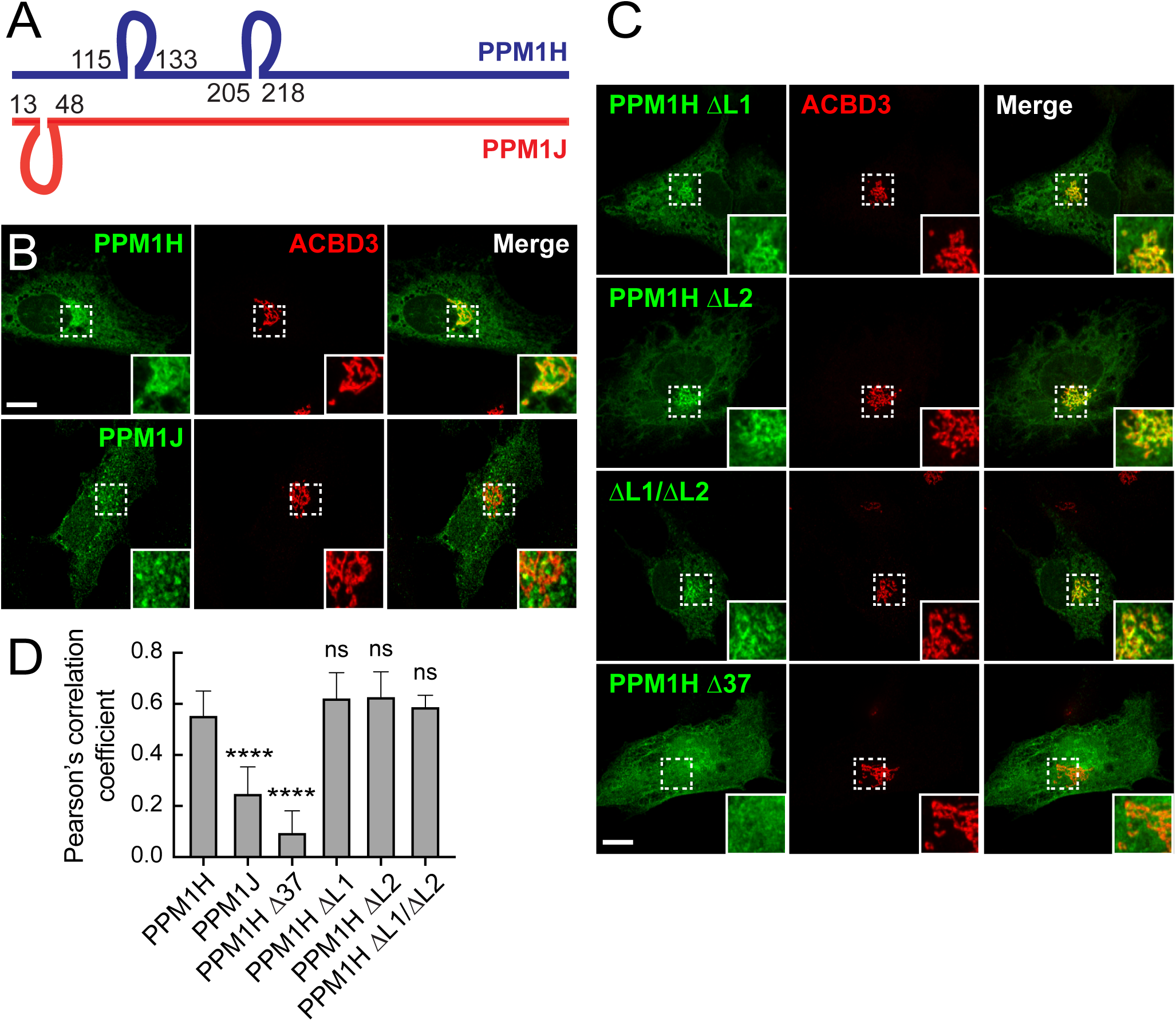
PPM1H is localized to the Golgi via its N-terminus. (A) Diagram of sequence alignment between PPM1H (blue) and PPM1J (red). (B, C) Immunofluorescence microscopy of RPE cells transiently transfected with plasmids encoding either HA-PPM1H, HA-PPM1J, HA-Δ37-PPM1H, HA-ΔLoop1-PPM1H (missing residues 115-133), HA-ΔLoop2-PPM1H (missing residues 205-218) or HA-ΔLoop1/2-PPM1H (missing both loops). After 24 hours cells were fixed and stained with mouse anti-HA antibody (green) and with rabbit anti-ACBD3 to label the Golgi (red). Scale bar, 10 µm. Shown are maximum intensity projections. Areas boxed with dashed lines are enlarged at lower right. (D) Co-localization of PPM1H or PPM1J with ACBD3 was determined by Pearson’s coefficient. Error bars represent SEM from two independent experiments. Significance was determined relative to HA-PPM1H by one way ANOVA. In relation to HA-PPM1H, p values were: ****p<0.0001 for HA-PPM1J, ****p<0.0001 for HA-Δ37 PPM1H, p=0.084373 for HA-ΔLoop1-PPM1H, p=0.060330 for HA-ΔLoop2-PPM1H, and p=0.421158 for HA-ΔLoop1/2-PPM1H.

### Golgi surveillance by PPM1H phosphatase

Phosphatase localization can play an important role in regulation of phospho-substrate selectivity. Rab8A, 10, 12, 29, 35 and 43 are all LRRK2 substrates and several of these are normally localized at or near the Golgi complex; Rab8A, Rab10 and Rab12 are the most predominant and ubiquitously expressed LRRK2 substrates (2,3). We determined the localizations of GFP-tagged Rab8A, 10, 12 and 29 in A549 cells, in relation to PPM1H-mApple. As shown in Figs. 2A,B, exogenous PPM1H showed the highest level of co-localization with Rab8A and Rab29; Rab10 and especially Rab12 showed less similar distributions but displayed some overlap with PPM1H. We showed previously strong co-localization of HA-PPM1H with exogenous GFP-Rab10 in RPE cells (11); the extent of co-localization is somewhat cell type specific. Thus, PPM1H on the Golgi appears to protect Golgi-associated Rab proteins from LRRK2 phosphorylation and inactivation; there is sufficient overlap for exogenously expressed, membrane-associated PPM1H to access each of its substrates, although they may be dephosphorylated with different efficiencies, with Rab12 showing the lowest extent of co-localization (Fig. 2B).

**Figure 2.**
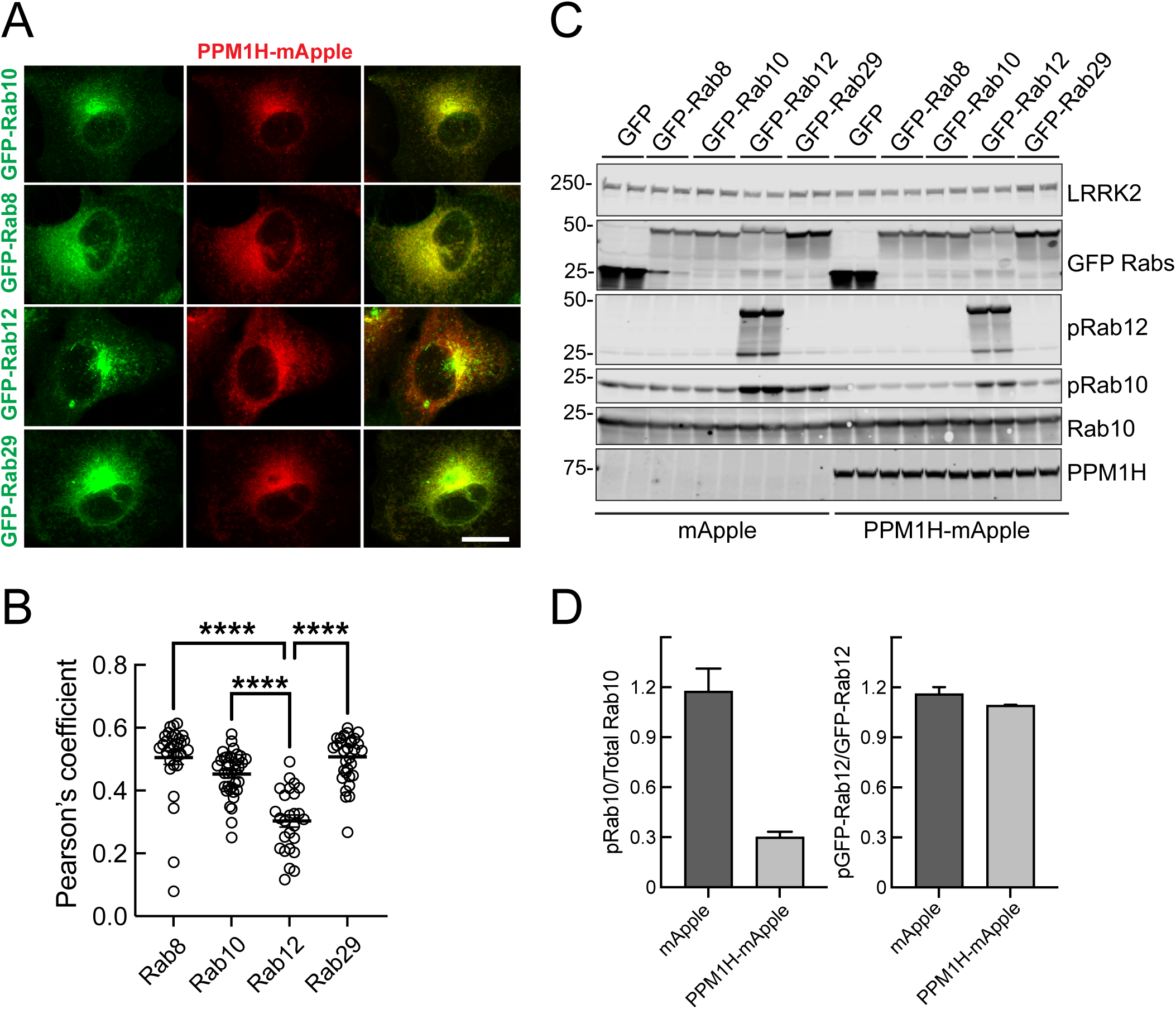
Localization of endogenous Rabs in A549 cells. (A) A549 cells were transduced with lentiviruses encoding PPM1H-mApple (red) and either GFP-Rab8A, GFP-Rab10, GFP-Rab12, or GFP-Rab29 (green) for stable overexpression. Scale bar, 10 µm. Representative images are shown as maximum intensity projections. (B) Co-localization of Rabs with PPM1H was determined by Pearson’s coefficient. Error bars represent SEM from two independent experiments with >20 cells per condition. Significance was determined relative to GFP-Rab12 by one way ANOVA. ****p<0.0001. (C) A549 cells transduced with lentiviruses as in A were lysed and analyzed by immunoblotting. Proteins were detected using mouse anti-LRRK2, chicken anti-GFP, mouse anti-RFP (for PPM1H), Rabbit anti-pRab12, mouse anti-total Rab10 and Rabbit anti-pRab10 antibodies. (D) Quantification of the ratio of pRab10 to total Rab10 (left) and pRab12 and total Rab12 (Right).

It is important to note that A549 cells contain only 37,000 molecules of PPM1H in relation to 2.6×10^6^ molecules of Rab10, 960,000 Rab8A, 135,000 Rab12, and 25,000 Rab29 molecules (https://copica.proteo.info/#/copybrowse/A549_single_shot.txt). Exogenously expressed PPM1H may show a broader distribution than the endogenous protein. In addition, we cannot distinguish whether the small percentage of a total Rab protein that is LRRK2 phosphorylated (10) has adequate access to endogenous PPM1H. Despite these limitations, Rab12 showed the least amount of co-localization with exogenously expressed PPM1H.

Immunoblot analysis of these cells confirmed that exogenous PPM1H was capable of dephosphorylating phosphoRab10 efficiently (Fig. 2C). However, phosphoRab12 was not efficiently dephosphorylated under these moderate expression conditions (Fig. 2C,D).

### PPM1H’s amphipathic helix is needed for Golgi localization and best activity

Analysis of the secondary structure of the PPM1H N-terminus revealed that it comprises an amphipathic helix, with Leu2, Val9 and Ile16 lying at the center of a possible hydrophobic face (Fig. 3A). This was reminiscent of work from Antonny and colleagues who showed that the Golgi-associated ArfGAP1 protein relies on an amphipathic helix to associate with the Golgi; this sequence is sufficient to drive liposome association and catalytic activation of ArfGAP1 protein (14,15).

**Figure 3.**
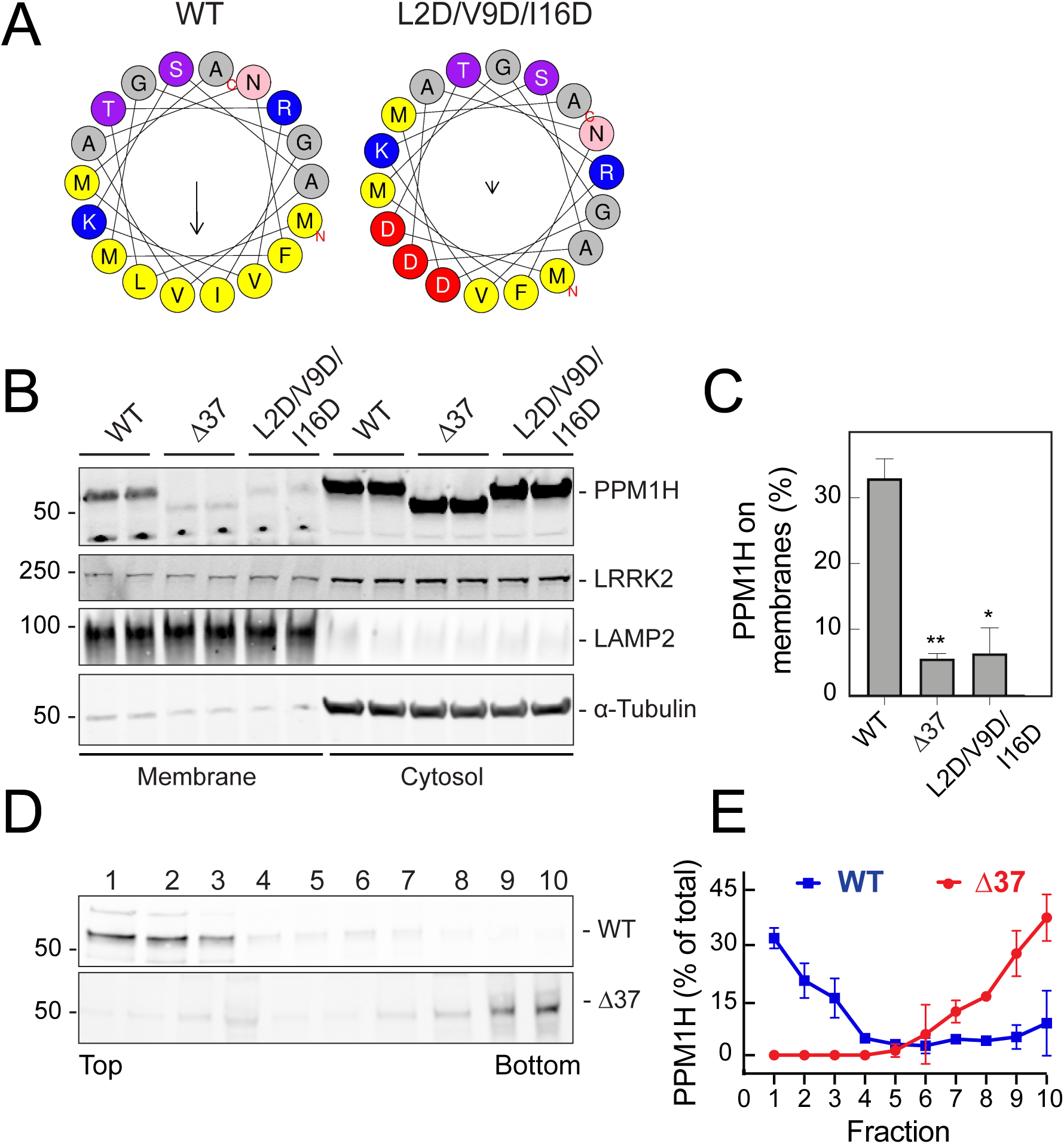
PPM1H’s N-terminal amphipathic helix is needed for membrane association. (A) Helical wheel projections made with Heliquest software (24) revealing the amphipathic helix at the N-terminus of PPM1H and location of L2D/V9D/I16D mutant residues introduced within the helix to block amphipathicity. (B,C) HEK293T cells were transiently transfected with plasmids encoding either HA-PPM1H, HA-Δ37-PPM1H or HA-L2D/V9D/I16D-PPM1H. After 36h cells were harvested and membrane and cytosol fractions obtained. 80 µg membrane protein and an equivalent volume of cytosolic fractions were processed for immunoblotting. Proteins were detected using mouse anti-LRRK2, mouse anti-LAMP2, mouse anti-HA, mouse anti alpha-tubulin antibodies. (C) Quantification of the fraction of HA-PPM1H on membranes. Error bars represent SEM from two independent experiments. Significance was determined relative to wtPPM1H by student’s T-test. **p=0.0056 for HA-Δ37-PPM1H and *p=0.0142 for HA-L2D/V9D/I16D-PPM1H. (D) Sucrose gradient flotation of wild type or Δ37 PPM1H in the presence of 30nm liposomes. The distribution of PPM1H across the gradient was determined by immunoblot; fractions were collected from the top. (E) Quantitation of PPM1H flotation from two independent experiments as in D.

To test the importance of the amphipathic nature of this sequence, we compared the subcellular fractionation properties of wild type, Δ37, and L2D/V9D/I16D proteins. Upon exogenous PPM1H expression and differential centrifugation to separate cytosolic from membrane associated proteins, ∼33% of PPM1H co-sedimented with membranes, in contrast with only ∼6% of Δ37 or L2D/V9D/I16D mutant proteins (Fig. 3B,C). For comparison, under these conditions, ∼10% of LRRK2 was membrane associated (8). These experiments demonstrate the importance of PPM1H’s N-terminal 37 residues for membrane association in cells, and show that residues 2, 9, and 16 are critical for membrane association–consistent with their role in stabilizing an amphipathic helix.

Analogous to ArfGAP1, PPM1H’s N-terminus was sufficient to enable purified PPM1H to associate with liposomes as monitored by sucrose gradient flotation (Figure 3D,E). In these experiments, purified PPM1H or Δ37 PPM1H were incubated together with 30nm liposomes (of Golgi lipid composition), overlaid with sucrose, and then ultracentrifuged to equilibrium. Full length PPM1H floated to the top of the gradient in the presence of liposomes while Δ37 PPM1H did not. This confirms that the N-terminal amphipathic helix drives PPM1H membrane association in vitro and in cells.

### Membrane associated PPM1H is most active

As might be expected for an enzyme that acts on membrane associated phosphoRab GTPases, full length PPM1H enzyme showed greatest activity in cells compared with truncated forms of the enzyme that lack 10, 20, 34 or 37 N-terminal residues of the amphipathic helix (Figure 4). These residues are predicted to be far from the PPM1H active site and were not present in the portion of PPM1H that was crystallized by Waschbüsch et al. (16). The activity of various PPM1H protein constructs was assessed by co-expression with hyperactive, Flag-tagged LRRK2 R1441C followed by immunoblotting to monitor decreases in phosphoRab10 levels (Fig. 4).

**Figure 4.**
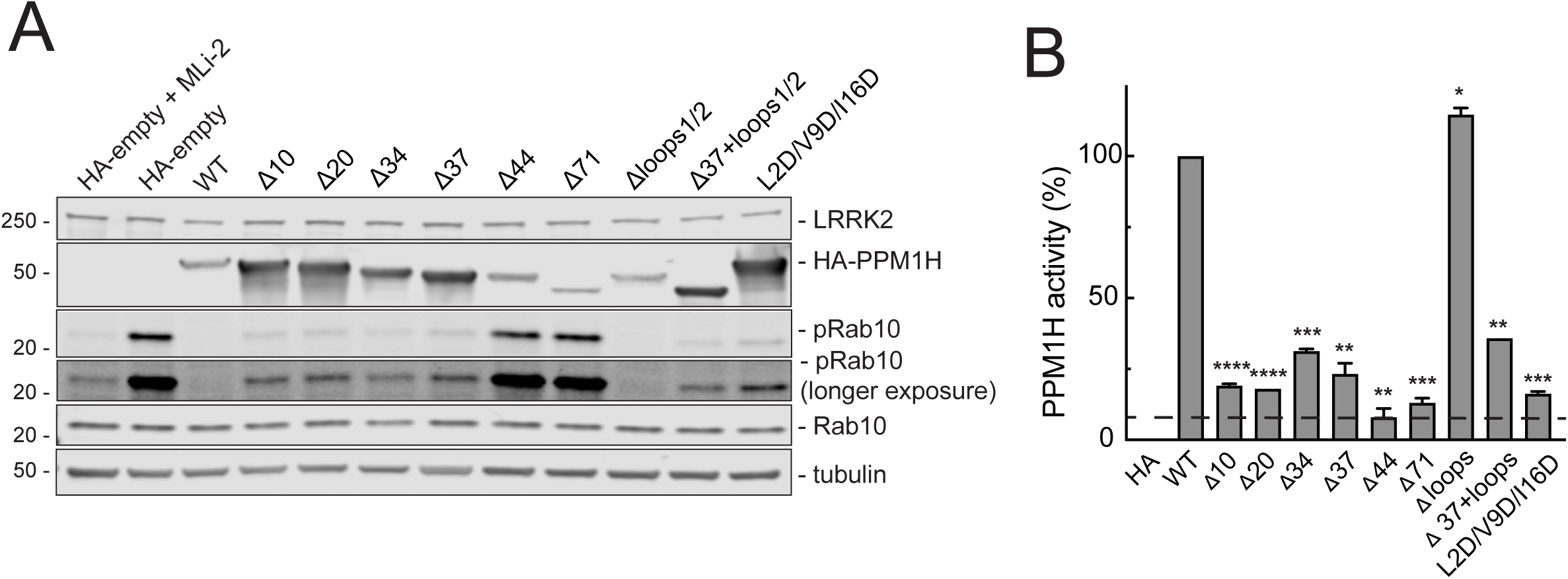
PPM1H’s amphipathic helix is needed for full activity in cells. (A) HEK293T cells were transiently transfected for 24h with Flag-LRRK2 R1441C together with either HA-empty, HA-PPM1H, HA-Δ10-PPM1H, HA-Δ20-PPM1H, HA-Δ34-PPM1H, HA-Δ37-PPM1H, HA-Δ44-PPM1H, HA-Δ71-PPM1H, HA-ΔLoop1/2-PPM1H, HA-Δ37/ΔLoop1/2-PPM1H or HA-L2D/V9D/I16D-PPM1H. After 24 hours cells were lysed and 30 µg protein analyzed by immunoblotting. Proteins were detected using mouse anti-LRRK2, rabbit anti-HA, rabbit anti-pRab10, mouse anti-total Rab10 and mouse anti alpha-tubulin antibodies. (B) Activity of PPM1H was quantified by normalizing pRab10/total Rab10 levels relative to PPM1H expression, with WT PPM1H set at 100% activity. Error bars represent SEM from two independent experiments. Significance was determined relative to HA-PPM1H by one-way ANOVA; ****P<0.0001 for HA-Δ10-PPM1H, HA-Δ20-PPM1H, ***P=0.0001 for HA-Δ34-PPM1H and HA-L2D/V9D/I16D-PPM1H, **P=0.0024 for HA-Δ37-PPM1H, **P=0.0012 for HA-Δ44-PPM1H, ***P=0.0004 for HA-Δ71-PPM1H, *P=0.0255 for HA-ΔLoop1/2-PPM1H, **P=0.0064 for HA-Δ37/ΔLoop1/2-PPM1H.

As expected, wild type PPM1H expression abolished detectable phosphoRab10 (Fig. 4A, lane 3). When normalized to the expression level of each PPM1H construct, truncations of 10, 20, 34 or 37 residues greatly decreased PPM1H activity, at least upon overexpression in HEK 293T cells (Fig. 4B). Longer N-terminal truncations of 44 or 71 residues completely blocked PPM1H activity, confirming prior work from Waschbüsch et al. (2021). Loss of the amphipathic helix by insertion of charged residues (L2D/V9D/I16D) into the predicted face of the helix also eliminated most PPM1H activity in this overexpression paradigm (Fig. 4A,B). No activity decrease was seen for PPM1H in which the loops at positions 115-133 and 205-218 (loops1/2) were deleted (Fig. 4). Mutation of the potential S124 and S211 phosphorylation sites was also without consequence (Supplemental Figure 3).

These experiments suggest that membrane association increases PPM1H’s activity in cells. Consistent with this possibility, highly curved liposomes directly activated purified PPM1H enzyme in vitro. Shown in Figure 5 is the loss of phosphoRab10 protein as a function of time in the presence of PPM1H protein. Inclusion of highly curved, small liposomes (30nm) but not larger, 100nm liposomes, activated PPM1H activity in these in vitro reactions (Fig. 5B). These experiments were carried out with extremely limiting levels of PPM1H protein (15 ng) in reactions containing much larger amounts of in vitro phosphorylated Rab10 protein (1.5µg).

**Figure 5.**
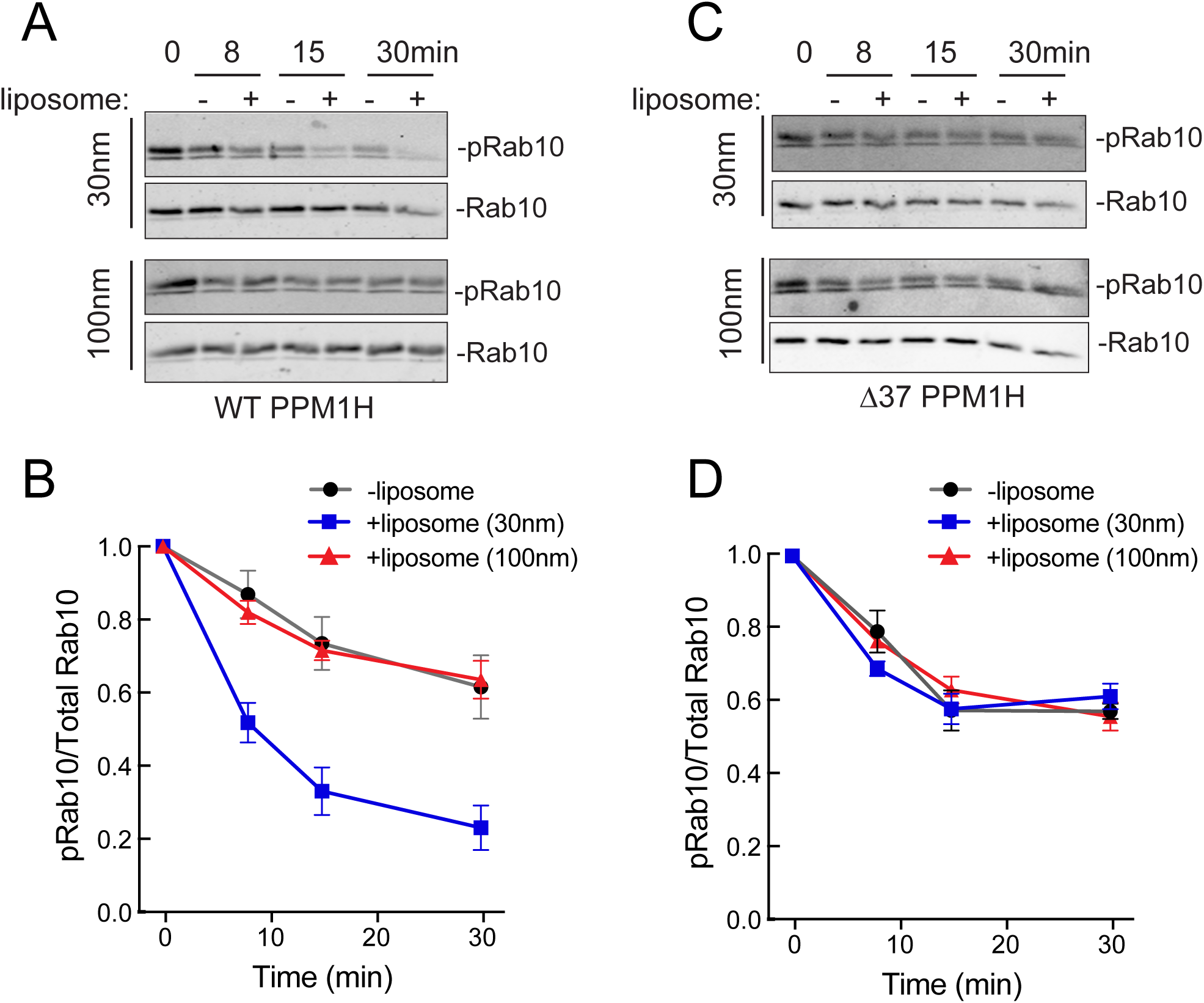
Activation of PPM1H by highly curved membranes requires its amphipathic helix. Purified PPM1H (A) or Δ37-PPM1H (C) proteins were assayed for phosphatase activity in the presence of 30nm or 100nm liposomes for the times indicated_[CC1]_. The respective rates of decrease of phosphoRab10 in each reaction was used to monitor phosphatase activity (B,D). Error bars represent SEM from 2 independent experiments carried out in duplicate.

Importantly, the activity of Δ37 PPM1H was completely unaffected by the presence of small or large liposomes (Fig. 5C,D). This suggests that interaction of PPM1H’s N-terminus with a highly curved membrane in some way alters its structure to activate its catalytic activity. In the crystal structure of PPM1H missing residues 1-32 (16), N-terminal residues 33-79 follow an irregular path behind the active site that spans two beta sheets of the catalytic domain. Perhaps highly curved membrane vesicle association tugs on this portion of the enzyme to orient the substrate for most efficient catalysis. These data support the conclusion that membrane associated PPM1H is more active than cytosolic PPM1H protein.

### Relocalization of PPM1H reveals its potency at the mother centriole

A phosphatase that is partly cytosolic should theoretically have access to Rab GTPases wherever they are located in cells. However, if membrane localization is truly important for substrate access or activation, artificial anchoring of PPM1H in different locations should provide important clues to the significance of its endogenous localization. Thus, we relocalized PPM1H to distinct cellular compartments and monitored the consequences for phosphoRab10 levels and for primary cilia formation that is a highly sensitive measure of PPM1H activity (11).

PPM1H was first re-localized to mitochondria by attaching an amphipathic helix derived from monoamine oxidase (17) to the PPM1H C-terminus; this hybrid protein will be referred to as mito-PPM1H. Note that the mitochondrial targeting signal dominated the targeting process when added to either full length PPM1H protein or Δ37 PPM1H (Figure 6A).

**Figure 6.**
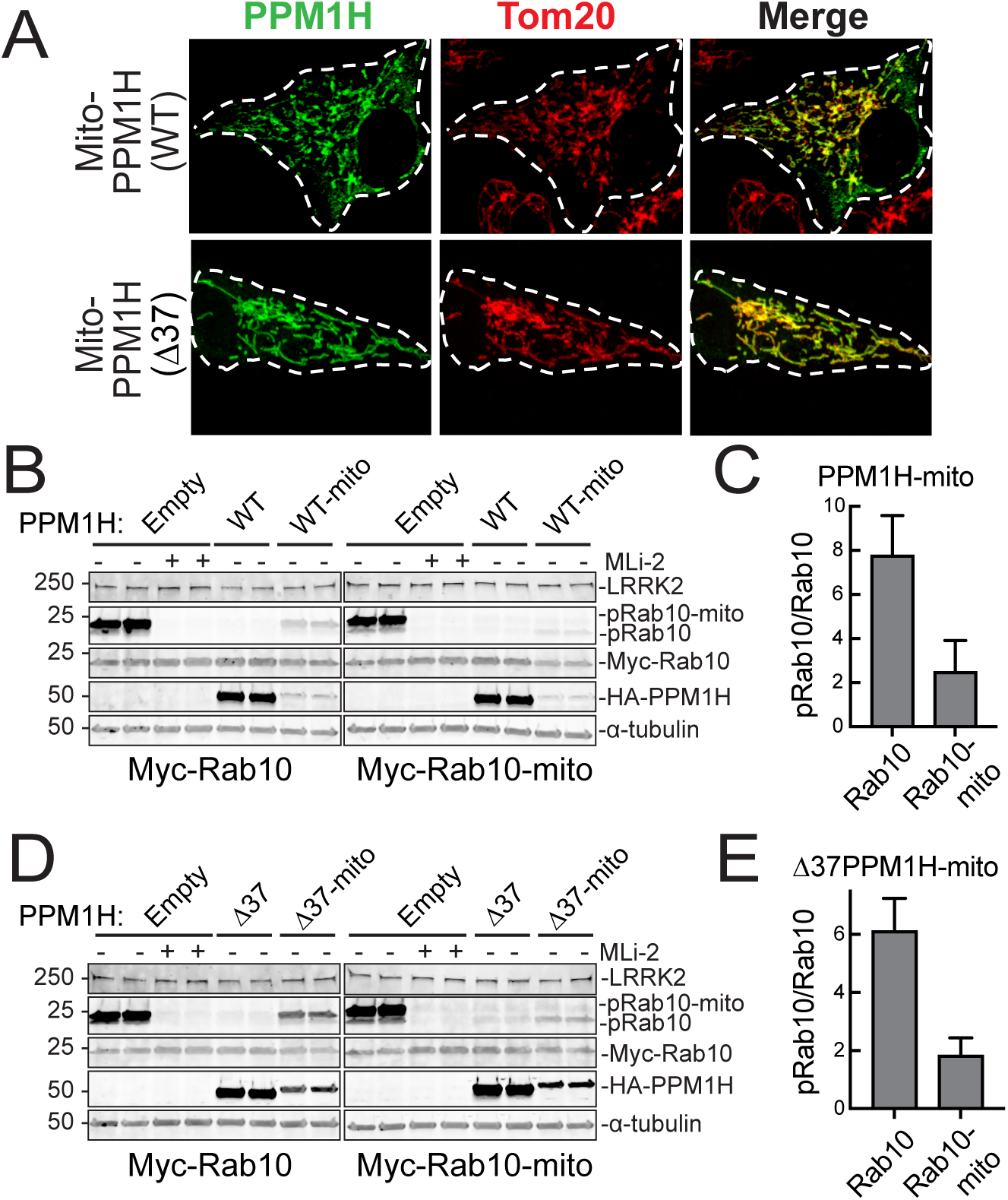
Mitochondrially localized PPM1H is catalytically active. (A) Representative localizations of PPM1H-mito and Δ37-PPM1H-mito proteins in RPE cells. Magnification bar, 10µm. (B,C) Dephosphorylation of wild type or mitochondrially targeted Myc-Rab10 by wild type or mitochondrially targeted PPM1H in HEK293T cells as indicated. Quantitation shown in C and E represent the mean of two independent experiments. Error bars represent SEM. (D,E) Same as in B,C using Δ37PPM1H or Δ37PPM1H-mito tag. Quantitation of changes in mito-pRab10 was carried out by analyzing only the upper mito-pRab10 band.

We next explored the ability of wild type or mitochondrially anchored PPM1H to dephosphorylate Rab10 or mitochondrially anchored Rab10. In these experiments, the PPM1H was much more highly expressed than the mitochondrially anchored protein (Fig. 6B,D) but comparisons could nevertheless be made between those amounts of mito-PPM1H acting on either wild type or mitochondrially anchored Rab10 proteins. Not surprisingly, mitochondrially anchored PPM1H proteins (mito-PPM1H or Δ37 mito-PPM1H) were better able to dephosphorylate mitochondrially anchored Rab10 protein than wild type Rab10 (Fig. 6 C,E). Wild type (full length, Fig. 6B) and Δ37 PPM1H (Fig. 6D) efficiently dephosphorylated wild type and mitochondrially anchored Rab10 at these expression levels, presumably due to diffusion. These experiments confirm that catalytic activity of mitochondrially localized PPM1H forms.

We also anchored PPM1H stably on the Golgi complex and at the mother centriole by appending sequences from TMEM115 (Golgi) or the PACT domain (centriole). As shown in Figure 7A, Δ37 PPM1H represented the cytosolic form; Δ37 TMEM115 PPM1H was exclusively Golgi localized, Δ37 PPM1H PACT was mostly centriolar but some cells also showed a broader cytoplasmic distribution; Δ37 PPM1H-mito was constrained to mitochondria. The expression of each of these constructs was under Doxycycline control, and addition of doxycycline turned on the expression of proteins of the correct molecular mass (Fig. 7B).

**Figure 7.**
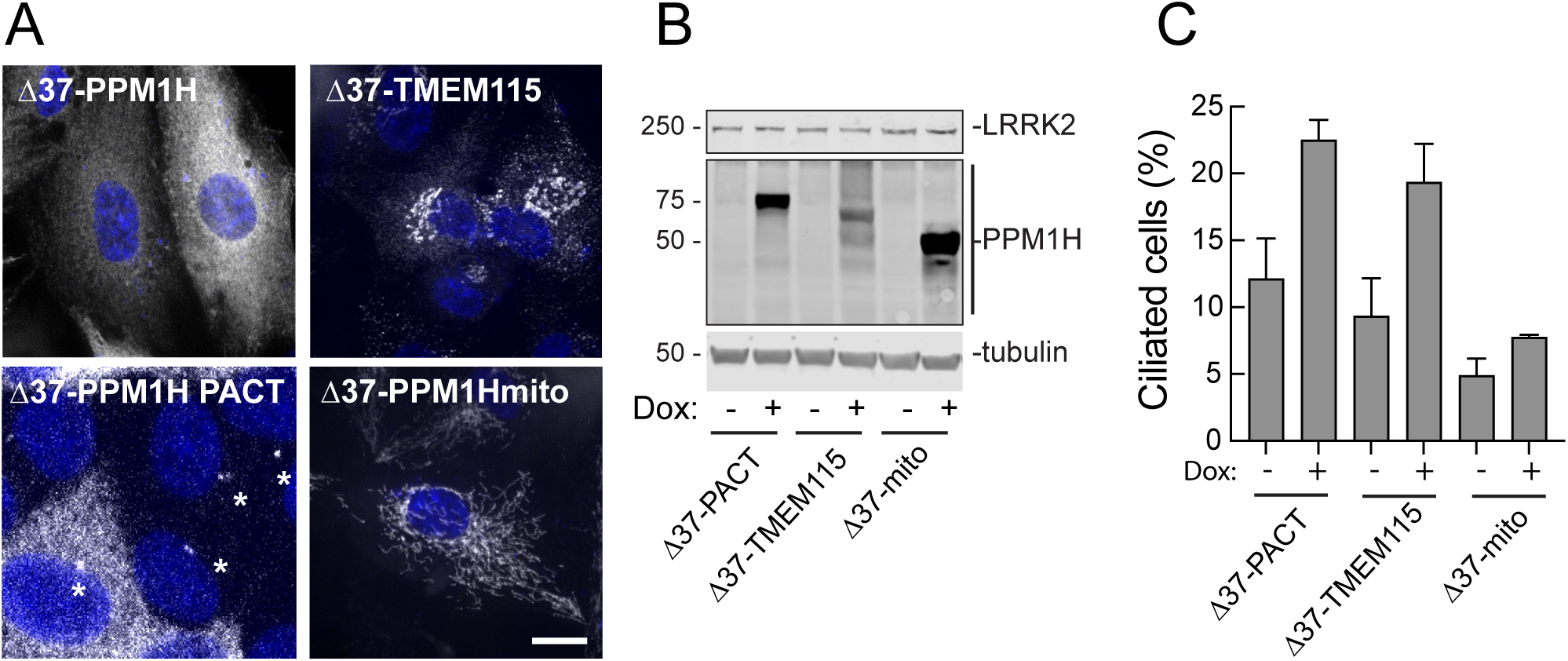
PPM1H influences ciliogenesis from the mother centriole or Golgi but not mitochondria. (A) PPM1H knockout A549 cells were transduced with lentiviruses encoding HA-tagged Dox-inducible Δ37-PPM1H, Δ37-PPM1H-PACT, Δ37-PPM1H-TMEM115 and Δ37-PPM1H-mito. HA-PPM1H signal was detected by using mouse anti-HA antibody. Shown are the localizations of indicated constructs. Magnification bar, 10µm. (B) Immunoblot of construct expression for the experiments presented in A. Proteins were detected using mouse anti-LRRK2, mouse anti-HA and mouse anti alpha-tubulin antibodies. (C) Percent ciliated cells with or without addition of Doxycycline (DOX) to induce expression of the indicated constructs. Shown is the mean of two independent experiments. For the PACT construct, only cells displaying uniquely centrosomal PPM1H localization were scored. Error bars represent SEM from two independent experiments with >300 cells per condition.

Using this sensitive and controlled expression system, Golgi and centriole-anchored PPM1H were both able to greatly stimulate the percentage of A549 cells that were ciliated, presumably due to decreased phosphoRab10 levels. This was in contrast with cells expressing much higher levels of mito-PPM1H. This experiment shows that PPM1H localized to the centriole or Golgi region can access and act upon phosphoRabs to permit primary cilia formation.

### PPM1H acts on phosphoRab12 in vitro but not in vivo

As mentioned above, previous work showed that phosphoRab8A and phosphoRab10 are much better PPM1H substrates than phosphoRab12 in cells (Berndsen et al., 2019; see also Fig. 2). Moreover, a PPM1H substrate-trapping mutant failed to trap Rab12 in cells as determined by mass spectrometry (Berndsen et al., 2019). Thus, in vitro dephosphorylation experiments have focused on phosphoRab8A and phosphoRab10 (11.16). Given the proportionally lower extent of co-localization of Rab12 with PPM1H in relation to Rab8A and Rab10, it was possible that phosphoRab12 is a much poorer substrate for PPM1H in cells simply because the two proteins co-localize less well (Figure 2). Alternatively, PPM1H enzyme may discriminate between Rab substrates and simply prefer Rab8A and Rab10.

We tested the ability of purified PPM1H to dephosphorylate phosphoRab10 and phosphoRab12 to distinguish between these possibilities. Figure 8 shows the kinetics of PPM1H-mediated Rab10 and Rab12 dephosphorylation: both Rabs were dephosphorylated at the same rate in vitro when assayed in parallel with limiting amounts of purified enzyme. Thus substrate selectivity cannot explain the inefficient dephosphorylation of phosphoRab12 in cells.

**Figure 8.**
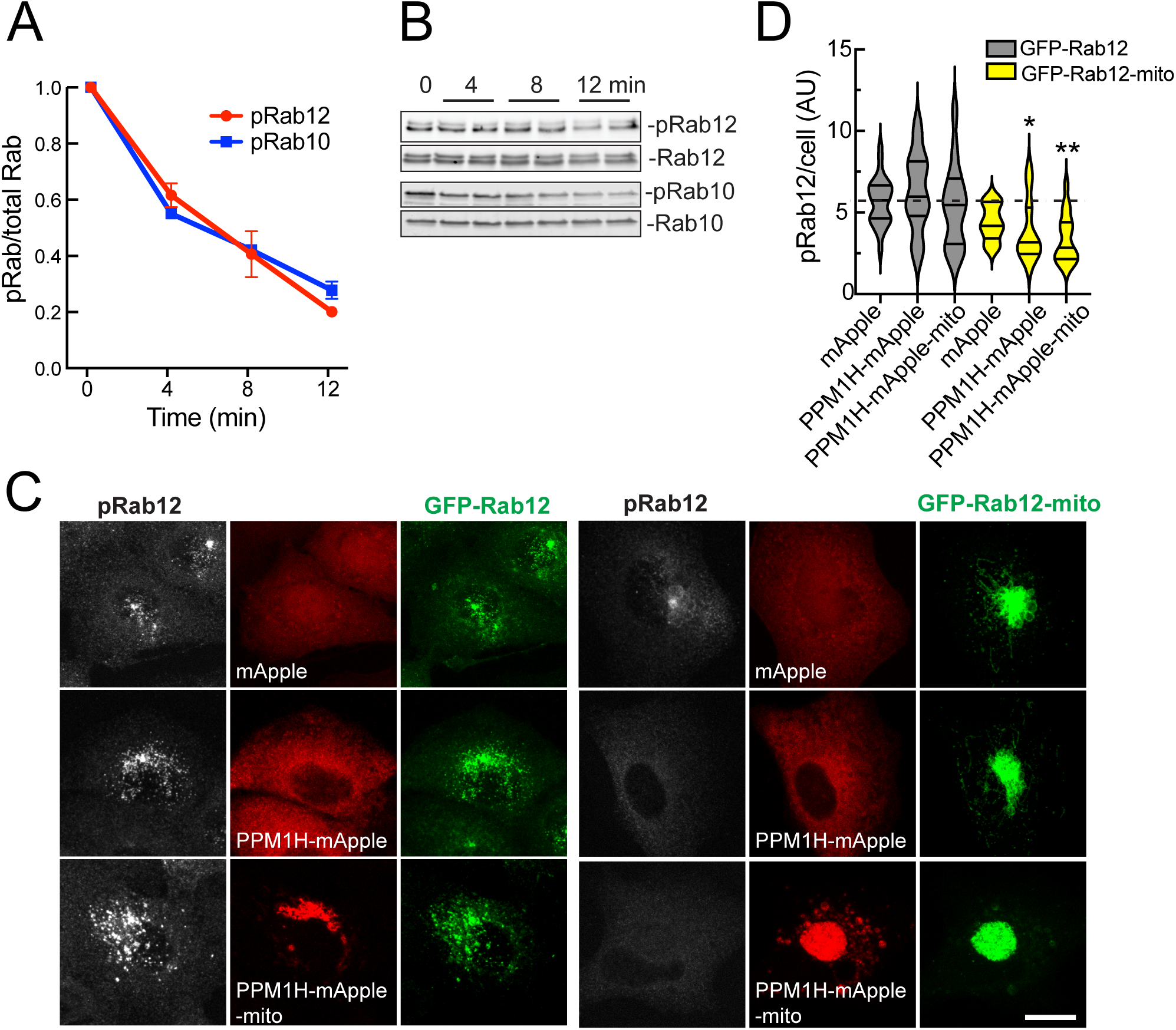
Localization is an important determinant of pRab12 dephosphorylation by PPM1H. (A) Purified PPM1H was assayed for phosphatase activity with purified pRab12 or pRab10 in the presence of 30nm liposomes for the times indicated. Phosphatase activity was monitored by immunoblot (as in B) to detect loss of pRab12 or pRab10 as a function of time. Shown are the means of 2 independent experiments carried out in duplicate. (C) A549 cells were transduced with lentiviruses encoding mApple (red), PPM1H-mApple or PPM1H-mApple-mito and either GFP-Rab12 (green) or GFP-Rab12-mito to create stably expressing cell lines. Cells were subsequently stained with anti-pRab12 antibody (white). Scale bar, 10 µm. Representative images are shown as maximum intensity projections. (D) Shown is the amount of pRab12 per cell as quantified as a mean integrated density. Error bars represent SEM from three independent experiments with at least 30 cells per condition. Significance was determined relative to values in GFP-Rab12-mito cells expressing mApple by T-test; *P=0.0424 for PPM1H-mApple, *P=0.0369 for PPM1H-mApple-mito.

To further evaluate the importance of localization on PPM1H substrate selectivity, we re-localized both Rab12 and PPM1H to mitochondria and monitored the ability of PPM1H to dephosphorylate Rab12. Figure 8C (left panels) shows the localizations of GFP-Rab12, phosphoRab12 and PPM1H in cells expressing mApple, PPM1H mApple, or PPM1H mApple that is mitochondrially anchored. In all cases, phosphoRab12 levels were unchanged (Figure 8C,D). In contrast, when Rab12 was localized to mitochondria, it was LRRK2 phosphorylated to a small extent, and was a much better substrate for mitochondrially anchored PPM1H enzyme Fig. 8C, D). Wild type PPM1H also acted a little bit on mitochondrially anchored Rab12 protein; we believe this is most likely due to the small amount of mitochondrially localized wild type PPM1H protein (Supplemental Figure 1). These experiments strongly suggest that Rab12 localization explains why it is a poor PPM1H substrate relative to Rab8A and Rab10 proteins in cells; if it was somehow protected from PPM1H action due to effector binding, that effector should have been able to access phosphoRab12 on mitochondrial surfaces. Indeed, mitochondrially targeted Rab12 causes collapse of the mitochondria into a bundle; this is likely due to an effector interaction but it does not impede PPM1H action.

### Summary

We have shown here that PPM1H is a predominantly Golgi-localized enzyme with high colocalization with Rab8A, Rab10, and Rab29 in A549 cells and with GFP-Rab10 in RPE cells. This localization is consistent with PPM1H’s role as a phosphoRab GTPase-specific phosphatase (11). Membrane association of PPM1H in cells and in vitro is driven by the protein’s N-terminal 37 residues that are predicted to form an amphipathic helix. Indeed, these residues drive liposome association with preference for small liposomes that have a very high degree of membrane curvature. This is reminiscent of ArfGAP1, a Golgi associated enzyme that is important for COP-I vesicle formation and is activated on highly curved membranes via an internal amphipathic helix (14,15).

High membrane curvature occurs at the rims of the Golgi complex and proteins containing amphipathic helices may accumulate in that sub-domain. Importantly, the ciliary pocket at the base of the primary cilium is also highly curved and an intriguing possibility is that endogenous PPM1H becomes concentrated there. Unfortunately it is impossible to determine the localization of endogenous PPM1H protein as it occurs in cells at ∼37000 copies in relation to millions of Rab8A and Rab10 proteins and could not be reliably detected by a recently available commercial antibody (AbCam EPR26028-53) when knockout cells were compared. Nevertheless, siRNA depletion of the few copies of PPM1H in wild type mouse embryonic fibroblasts is sufficient to block ciliogenesis, demonstrating a normal role of LRRK2-mediated Rab phosphorylation in regulating ciliogenesis (11).

PPM1H is likely to work in proximity with LRRK2 on membrane surfaces, as Rab phosphorylation shows rapid turnover and is rapidly reversed upon addition of kinase inhibitors (10,11). In this regard, it is interesting to consider the possibility that lysosome-associated LRRK2 hyperactivation upon lysosomal membrane damage (18,19) is exacerbated by the possible absence of PPM1H at that location. In addition, PPM1H alone cannot explain reversal of Rab GTPase phosphorylation: phosphoRab12 for example, is a poor PPM1H substrate and is likely a substrate for another phosphatase. Future work will elaborate the full panel of phosphatases that balance the action of LRRK2 kinase on Rab GTPase phosphorylation.

## Key Resources Table

| Key Resources Table |  |  |  |  |
| --- | --- | --- | --- | --- |
| Reagent type (species) or resource | Designation | Source or reference | Identifiers | Additional information |
| Antibody | anti-LRRK2 (mouse monoclonal) | Neuromab | N241A/34 | (1:1000) |
| Antibody | anti-Rab10 (mouse monoclonal) | Nanotools | 0680–100/Rab10-605B11 | (1:1000) |
| Antibody | anti-Rab10 (phospho T73) (rabbit monoclonal) | Abcam | ab230261 | (1:1000) |
| Antibody | anti-PPM1H (Sheep polyclonal) | MRC-PPU | F-1804 | (1:1000) |
| bacterial strain | E. coli DH5α | Thermo Fisher | 18258012 | |
| bacterial strain | E. coli STBL3 | Thermo Fisher | C737303 |  |
| bacterial strain | E. coli Rosetta DE3 pLys | Millipore | 70956 |  |
| cell line | RPE | ATCC | CRL-2302 |  |
| cell line | HEK293T | ATCC | CRL-3216 |  |
| Chemical compound, drug | MLi-2 | MRC PPU |  |  |
| Filter | Drain disc 10mm | Sigma Aldrich | WHA230300 |  |
| Filter | Polycarbonate membranes 0.05mm | Avanti Polar Lipids Inc. | 610003-1EA |  |
| Filter | Polycarbonate membranes 0.1mm | Avanti Polar Lipids Inc. | 610005-1EA |  |
| Filter | Polycarbonate membranes 0.4mm | Avanti Polar Lipids Inc. | 610007-1EA |  |
| Instrument | hand held mini extruder with heating block | Avanti Polar Lipids Inc. | 610000-1EA |  |
| Lipid | 18:1 (Δ9-Cis) PC (DOPC) | Avanti Polar Lipids | #850375 | 11 μmol |
| Lipid | 18:1 PS (DOPS) | Avanti Polar Lipids | #840035 | 5 μmol |
| Lipid | 18:1 PI(4)P | Avanti Polar Lipids | #850151 | 0.15 μmol |
| plasmid | pcDNA 3.1 myc-Rab10 | Addgene in progress |  | human |
| plasmid | pcDNA 3.1 myc-Rab10-mito | Addgene in progress |  | human |
| plasmid | pcDNA5D HA-PPM1H | MRC PPU | DU62789 | human |
| plasmid | pcDNA5D HA-Δ10 PPM1H | Addgene in progress | | human |
| plasmid | pcDNA5D HA-Δ20 PPM1H | Addgene in progress |  | human |
| plasmid | pcDNA5D HA-Δ34 PPM1H | Addgene in progress |  | human |
| plasmid | pcDNA5D HA-Δ37 PPM1H | Addgene in progress |  | human |
| plasmid | pcDNA5D HA-Δ44 PPM1H | Addgene in progress |  | human |
| plasmid | pcDNA5D HA-Δ71 PPM1H | Addgene in progress |  | human |
| plasmid | pcDNA5D HA-PPM1H AH mutant | Addgene in progress |  | human |
| plasmid | pcDNA5D HA-Δ115-133, 205-218 PPM1H | Addgene in progress |  | human |
| plasmid | pcDNA5D HA-Δ37, 115-133, 205-218 PPM1H | Addgene in progress |  | human |
| plasmid | pcDNA5D HA-PPM1J | MRC PPU | DU68077 | human |
| plasmid | pET15b+His-SUMO PPM1H | MRC PPU | DU62790 | human |
| plasmid | pET15b+His-SUMO Δ37 PPM1H | MRC PPU | DU68537 | human |
| plasmid | pET15b+His SUMO Rab10 (1-181) Q68L | Addgene in progress |  | human |
| Plasmid | CSII-PPM1H-mApple | Addgene |  | human |
| plasmid | pCMV5 FLAG-LRRK2 | MRC PPU | DU6841 | human |
| plasmid | pCMV5 Flag-LRRK2 R1441C | MRC PPU | DU13078 | human |
| Plasmid | pMCB306-GFP-Rab12 | Addgene | 198471 | human |
| Plasmid | pMCB306-GFP-Rab10 | Addgene | 130883 | human |
| Plasmid | pMCB306-GFP-Rab8 | Addgene | 198470 | **human** |
| Plasmid | pMCB306-GFP-Rab29 | Addgene | 198472 | **human** |
| Plasmid | pET 15b+His SUMO Rab12 (Q101L) | Addgene in process |  | human |
| Plasmid | Lenti-TO-HA-Δ37 PPM1H | Addgene in process |  | human |
| Plasmid | Lenti-TO-HA-Δ37 PPM1H -PACT | Addgene in process |  | human |
| Plasmid | Lenti-TO-HA-Δ37 PPM1H-TMEM115 | Addgene in process |  | human |
| Plasmid | Lenti-TO-HA-Δ37 PPM1H-mito | Addgene in process |  | human |
| Software, Algorithm | FIJI | PMID: 29187165 | RRID:SCR_002285 |  |
| Software, Algorithm | CellProfiler | PMID: 29969450 | RRID:SCR_007358 |  |
| Software, Algorithm | Chimera 2 | PMID:15264254 |  |  |

## Materials and Methods

### Cloning and plasmids

DNA constructs were amplified in E. coli DH5α or STBL3 and purified using mini prep columns (Econospin). DNA sequence verification was performed by Sequetech (http://www.sequetech.com). pET15b Rab10 Q68L 1-181 and pET15b His-Mst3 were kind gifts of Amir Khan (Harvard University). His-SUMO PPM1H full length, 38-514 were all cloned from pCMV5D HA-PPM1H into pET15b backbone.His SUMO Rab10(1-181) Q68L and His SUMO Rab12(Q101L) were cloned from pET15b Rab10 (1-181) Q68L and pQE 80L Rab12 (Q101L) respectively into pET15b backbone. pCMV5D HA-PPM1J (DU68077) and pCMV5D HA-PPM1H (DU62789) were obtained from the MRC-PPU Dundee. HA-PPM1H Δ10, Δ20, Δ34, Δ37, Δ44, Δ71 were all cloned from pCMV5D HA-PPM1H by PCR. PPM1H phosphorylation site mutants (S124A, S211A, S124A/S211A) were cloned from pCMV5D HA-PPM1H by site-directed mutagenesis using Q5 polymerase (New England Biolabs). To generate mitochondrially-targeted PPM1H, pCMV5D HA-PPM1H was first PCR amplified to contain an EcoRI site using 5’-GGGTAAGCGGCCGCTGATGACAGCTTGTTTCC-3’ and 5’-ACGCCTAAGAATTCGTCGAGTCTAGAGGGCCCGTT-3’ primers. A mitochondrial targeting sequence was PCR amplified from mito-Rab29 (5) using 5’-GGTGCGGCCGCCTTCTGGGAAAGGA-3’ and 5’-GAATAGGGGAATTCCCCTCAAGACCGTGGCAGGAG-3’ primers and added to the C-termini of HA-PPM1H or Δ37 HA-PPM1H between the NotI and EcoRI sites. To generate PPM1H-mApple-mito and GFP-Rab12-mito were generated by Gibson assembly in the CSII-PPM1H-mApple and pMCB306-GFP-Rab12 backbone (25).

### Cell culture and lysis

A549, RPE and HEK293T cells were purchased from ATCC. PPM1H knockout (KO) A549 cells were obtained from MRC-PPU. Cells were cultured in DMEM containing 10% (v/v) fetal calf serum, 2 mM L-glutamine, 100 U/ml penicillin, and 100 μg/ml streptomycin. All cells were grown at 37°C, 5% CO_2_ in a humidified atmosphere and regularly tested for Mycoplasma contamination. Unless otherwise indicated, cells were lysed in an ice-cold lysis buffer containing 50 mM Tris–HCl, pH 7.4, 1% (v/v) Triton X-100, 10% (v/v) glycerol, 0.15 M NaCl, 1 mM sodium orthovanadate, 50 mM NaF, 10 mM 2-glycerophosphate, 5 mM sodium pyrophosphate, 1 µg/ml microcystin-LR, and complete EDTA-free protease inhibitor cocktail (Roche). Lysates were clarified by centrifugation at 10,000g at 4°C for 10 min and supernatants protein was quantified by Bradford assay.

### Protein purification

His-SUMO-PPM1H and His-SUMO Δ37 PPM1H were purified according to this protocol <u>dx.doi.org/10.17504/protocols.io.bvjxn4pn</u>. The purified proteins were dialyzed overnight with Ulp1 SUMO protease (100ng/1mg of protein) to remove the His-SUMO tag from PPM1H. The dialyzed proteins were collected the next day and concentrated before application onto an S75 (24 ml) column fitted with 1ml HisTrap Ni-NTA column to remove the uncleaved His SUMO protein and the His-tagged Ulp1 SUMO protease.

### 293T overexpression assays

HEK293T cells were seeded into six-well plates and transiently transfected at 60-70% confluency using polyethylenimine (PEI) transfection reagent. 1 µg Flag-LRRK2 R1441C, 0.25 µg HA-PPM1H constructs and 6.25 µg PEI were diluted in 200 µL Opti-MEM™ Reduced serum medium (Gibco™) per well. 24 hours after transfection, cells were treated with 200 nM MLi-2 for 2 hours as indicated and lysed in an ice-cold lysis buffer. Samples were prepared for immunoblotting analysis as above.

### Immunofluorescence microscopy

A549 or RPE cells were transiently transfected with HA-PPM1H plasmids. After 24 h, cells were fixed with 4% (v/v) paraformaldehyde for 10 min, permeabilized with 0.1% Triton X-100 for 5 min, and blocked with 1% BSA for 1 h (dx.doi.org/10.17504/protocols.io.ewov1nmzkgr2/v1). Cells were subsequently stained with mouse or rabbit anti-HA antibody (Sigma-Aldrich H3663 or H6908, 1:1000) and the following Golgi markers: rabbit anti-GCC185 1:1000 (serum), rabbit anti-ß-Cop 1:1000 (serum), mouse anti-p115 1:1000 (ascites), mouse anti-cation independent mannose 6-phosphate receptor 1:1 (2G11 culture supernate), and rabbit anti-ACBD3 1:1000 (Sigma-Aldrich HPA015594). Highly cross absorbed H+L secondary antibodies (Life Technologies) conjugated to Alexa 488 or Alexa 568 were used at 1:5000. Primary and secondary antibody incubations were for 1 h at room temperature. Nuclei were stained with 0.1 µg/ml DAPI (Sigma). All images were obtained using a spinning disk confocal microscope (Yokogawa) with an electron multiplying charge coupled device (EMCCD) camera (Andor, UK) and a 100×1.4NA oil immersion objective. Images were analyzed using CellProfiler software (20,21) and presented as maximum intensity projections.

### Liposome preparation

Liposomes were generated with a composition of DOPC:DOPS:PI(4)P (69:30:1) (Avanti Polar Lipids; reference 22). A dried film of the indicated lipid mixture dissolved in chloroform was obtained by evaporation under a stream of nitrogen. The film was then resuspended in 50mM HEPES pH 7.5,120mM KCl. After two brief sonication cycles (5 sec each) using a bath sonicator, the liposome suspension was extruded 21 times sequentially through 0.4, 0.1 and 0.05 µm pore size polycarbonate filters using a hand extruder (Avanti). The final lipid concentration in the liposome suspension was 15mM.

### Sucrose density gradient flotation

Liposome flotation assays were performed following the method described by Bigay et al. (14) with slight modifications. Briefly, 1.5mM 30nm liposomes [DOPC:DOPS:PI(4)P] were incubated with 1mM PPM1H or Δ37-PPM1H for 15 min at 27°C in a total volume of 75μl in HKM buffer (20mM HEPES pH-7.5,150mM potassium acetate, 1mM magnesium chloride). 150μl of 75% w/v sucrose was then added and mixed to adjust the mixture to 50% sucrose. The high sucrose mixture was overlaid with 225μl, 25% w/v sucrose and 50μl of HKM buffer. The sample was centrifuged at 75,000 X g for 2.25 h in a Ti55 swinging bucket rotor. Ten 50μl fractions were manually collected using a pipetman by aspiration from the top of each tube. The fractions were then analyzed by immunoblotting to detect PPM1H or Δ37 PPM1H using sheep anti-PPM1H antibody.

### Immunoblot determination of membrane-associated PPM1H

HEK293T cells were transfected with either HA-PPM1H, HA-Δ37PPM1H or HA-L2D/V9D/I16DPPM1H constructs. 48 hrs after transfection, cells were washed 2x with ice cold PBS and swelled in 800µl of hypotonic buffer (10mM HEPES, PH=7.4). After 20 mins, 200µl of buffer (5x) was added to achieve a final concentration of 1x resuspension buffer (50 mM HEPES pH 7.4, 150 mM NaCl, 5 mM MgCl2, 0.5 mM DTT, 100 nM GDP, 1× protease inhibitor cocktail (Sigma) and 5 mM sodium fluoride, 2 mM sodium orthovanadate, 10 mM beta-glycerophosphate, 5 mM sodium pyrophosphate, 0.1 μg/ml Microcystin-LR). The suspension was passed 20 times through a 27G needle. Lysate was spun at 1000g for 5 mins to pellet nuclei. The post-nuclear supernatant was spun at 55,000 RPM for 20 min in a tabletop ultracentrifuge in TLA100.2 rotor; the resulting supernatant was collected as cytosolic fraction. Membrane pellet was solubilized in a resuspension buffer containing 1% Triton X-100. Protein concentrations were estimated by Bradford assay (Bio-Rad, Richmond, CA). Samples containing 80 μg of membrane protein, or the equivalent volume of cytosolic protein were processed for immunoblotting. All centrifugation steps were done at 4°C.

### PPM1H phosphatase assay in vitro

Untagged Rab10 (1-181) Q68L or untagged Rab12 (full length) Q101L was incubated with His-MST3 kinase in reaction buffer (50mM HEPES pH 8,100mM NaCl, 5mM MgCl_2_,100μM GTP, 0.5mM TCEP, 10% glycerol, 5μM BSA) at 4°C overnight to phosphorylate Rab10/Rab12 (For a detailed protocol see dx.doi.org/10.17504/protocols.io.bvjxn4pn). The next day, His-MST3 kinase was removed by passing the sample through a 1ml syringe column containing 100μl (50%) Ni-NTA slurry; the flow through containing phosphorylated Rab10/Rab12 was collected. 1.5µg phosphorylated Rab10/Rab12 was then incubated with 15ng PPM1H (For a detailed protocol see dx.doi.org/10.17504/protocols.io.bu7wnzpe) or 10ng Δ37 PPM1H in the presence or absence of liposomes at 30°C. Reactions were stopped by addition of 5X SDS-PAGE sample buffer. Samples were then analyzed by immunoblotting to detect for dephosphorylation of Rab10/Rab12 in the presence or absence of liposomes using anti-pRab10 (1:1000) antibody or anti-pRab12 (1:1000) antibody.

A detailed protocol for immunoblotting (23) is at https://dx.doi.org/10.17504/protocols.io.bsgrnbv6. 20µg protein was resolved by SDS PAGE and transferred onto nitrocellulose membranes using a Bio-Rad Trans-turbo blot system. Membranes were blocked with 2% BSA in Tris-buffered saline with Tween-20 for 30 min at room temperature (RT). Primary antibodies were diluted in blocking buffer as follows: mouse anti-LRRK2 N241A/34 (1:1000, Neuromab); mouse anti-Rab10 (1:1000, Nanotools); and rabbit anti-phospho Rab10 (1:1000, Abcam). Primary antibody incubations were overnight at 4°C. Li-COR secondary antibodies diluted in the blocking buffer were 680 nm donkey anti-rabbit (1:5,000) and 800 nm donkey anti-mouse (1:5000). Secondary antibody incubations were for 1h at RT. Blots were imaged using an Odyssey Infrared scanner (Li-COR) and quantified using ImageJ software.

### Ciliation

PPM1H KO A549 cells were infected with lentiviruses encoding indicated constructs and on day 3, infected cells were selected using 10µg/ml Blasticidin for 72hrs as described (dx.doi.org/10.17504/protocols.io.eq2ly7wpmlx9/v1); pools of stably infected cells were then assayed for cilia formation. Expression was induced using 1µg/ml doxycycline for 24 hours. Ciliation was monitored after 48 hr serum starvation using anti-Arl13B antibody (NeuroMab, Davis, California) to stain cilia for immunofluorescence microscopy (8).

## Acknowledgements

This study was funded by the joint efforts of The Michael J. Fox Foundation for Parkinson’s Research (MJFF) [MJFF-021132] and Aligning Science Across Parkinson’s (ASAP) initiative. MJFF administers the grant (ASAP 000463) on behalf of ASAP and itself.

For the purpose of open access, the authors have applied a CC-BY public copyright license to the Author Accepted Manuscript version arising from this submission. All primary data associated with each figure has been deposited in the Zenodo repository and can be found at https://doi.org/XXXXX/. We thank Kerryn Berndsen, Dario Alessi and Amir Khan for sharing data and helpful discussions.

---

**Supplemental figure 1.**
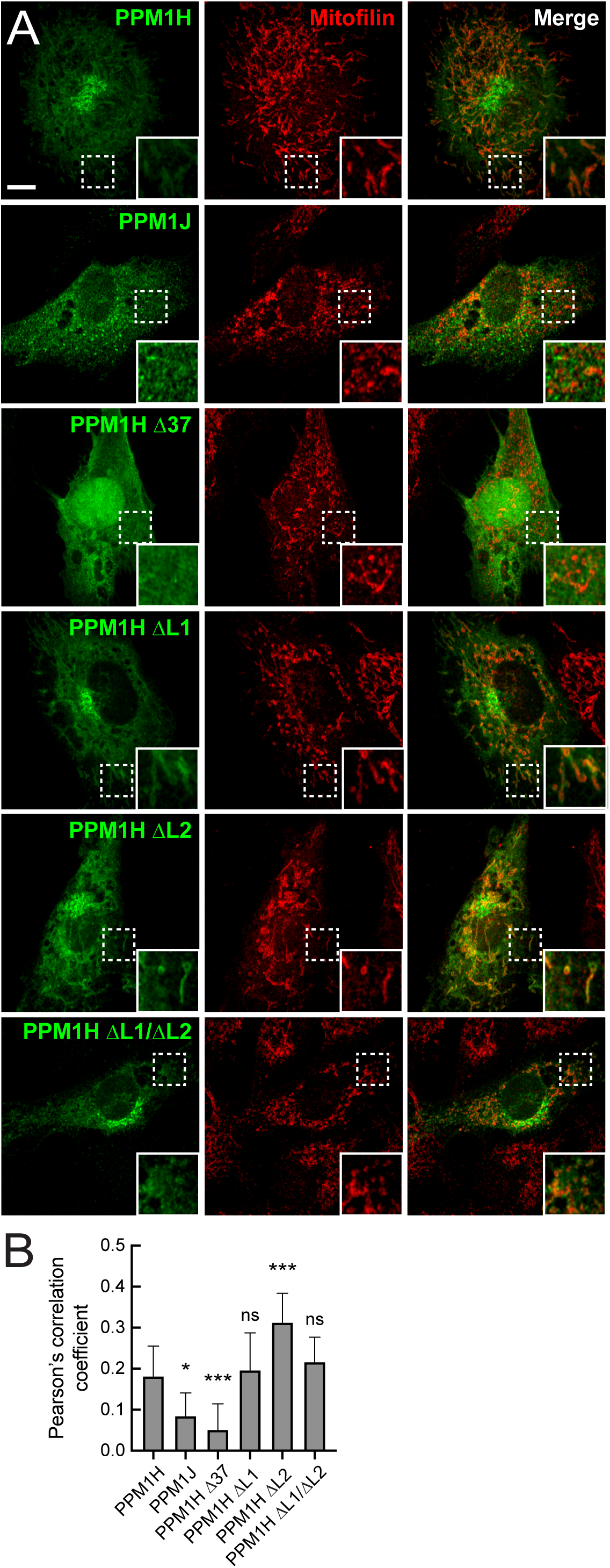
PPM1H constructs are mostly non-mitochondrial in RPE cells. RPE cells were transiently transfected with plasmids encoding either HA-PPM1H, HA-PPM1J, HA-Δ37-PPM1H, HA-ΔLoop1-PPM1H (missing 115-133), HA-ΔLoop2-PPM1H (missing 205-218) or HA-ΔLoop1/2-PPM1H (missing both loops). After 24 hours cells were subsequently fixed and stained with mouse anti-HA antibody (green) and rabbit anti-mitofilin (red). Scale bar, 10 µm. Shown are maximum intensity projections. Areas boxed with dashed lines are enlarged at lower right. (B) Co-localization of PPM1H or PPM1J with mitofilin was determined by Pearson’s coefficient. Error bars represent SEM from two independent experiments. Significance was determined relative to HA-wtPPM1H by one way ANOVA. *p=0.010169 for HA-wtPPM1H and HA-wtPPM1J, ***p=0.000339 for HA-wtPPM1H and HA-Δ37PPM1H, p=0.986811 for HA-ΔLoop1PPM1H, ***p=0.000202 for HA-wtPPM1H and HA-ΔLoop2PPM1H, p=0.628636 for HA-wtPPM1H and HA-ΔLoop1/2PPM1H.

**Supplemental figure 2.**
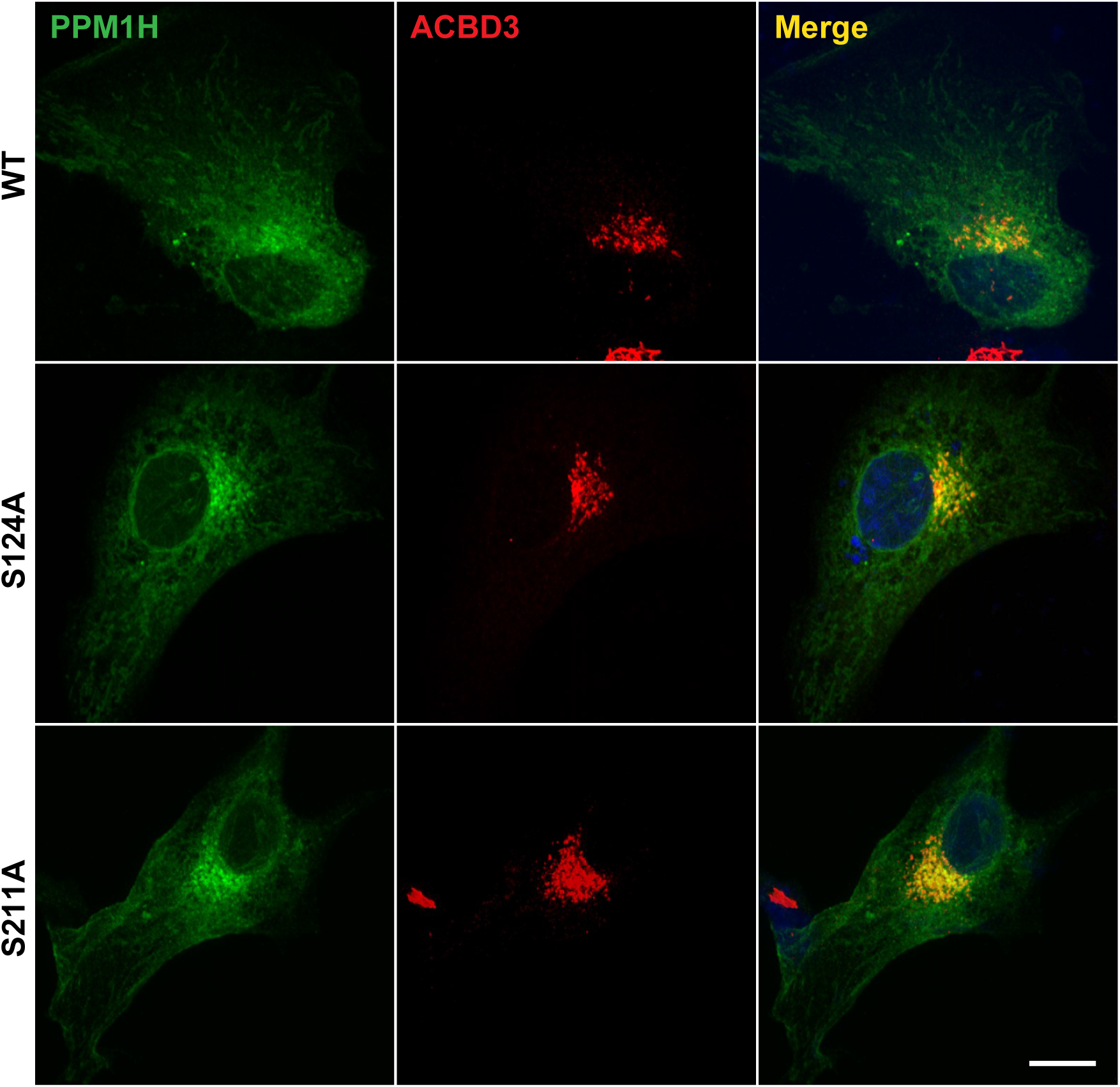
PPM1H phosphorylation mutants localize to the Golgi complex. The indicated PPM1H constructs were transfected and evaluated as in Figure 1. Scale bar, 10µm.

**Supplemental figure 3.**
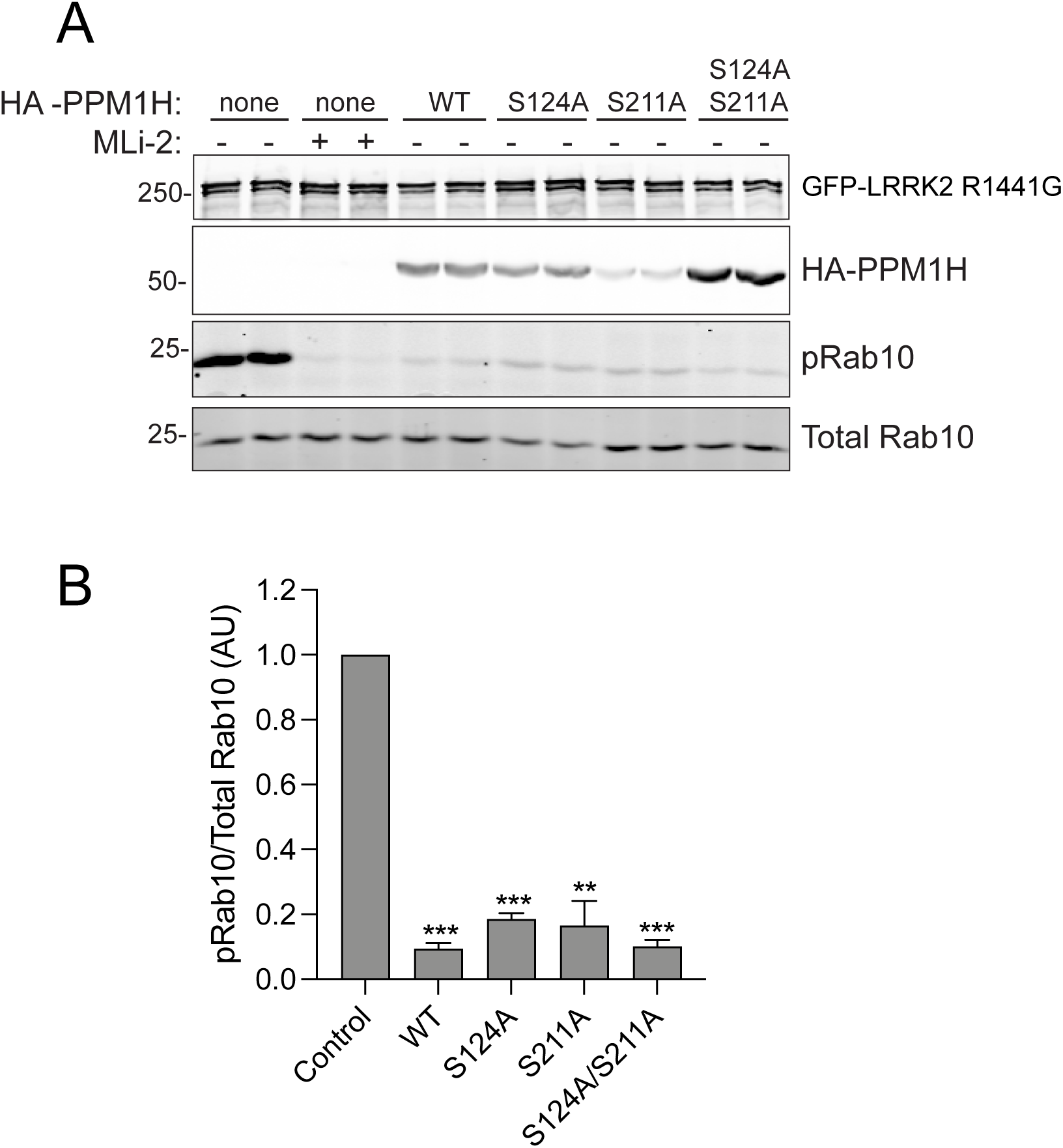
PPM1H phosphorylation site mutants are fully active. (A) HEK293T cells were transiently transfected for 24h with GFP-LRRK2 (R1441G) together with either HA-empty, HA-PPM1H, or HA-PPM1H S124A, S211A, or S124A/S211A. After 24 h cells were lysed and 30 µg protein analyzed by immunoblotting. Proteins were detected using mouse anti-LRRK2, rabbit anti-HA, rabbit anti-pRab10, and mouse anti-total Rab10 antibodies. (B) Activity of PPM1H was quantified as the ratio of pRab10 to total rab10. Error bars represent SEM from two independent experiments.

